# Gene regulatory network inference using mixed-norms regularized multivariate model with covariance selection

**DOI:** 10.1101/2022.12.21.521538

**Authors:** Alain J. Mbebi, Zoran Nikoloski

## Abstract

Despite extensive research efforts, reconstruction of gene regulatory networks (GRNs) from transcriptomics data remains a pressing challenge in systems biology. While non-linear approaches for reconstruction of GRNs show improved performance over simpler alternatives, we do not yet have understanding if joint modelling of multiple target genes may improve performance, even under linearity assumptions. To address this problem, we propose two novel approaches that cast the GRN reconstruction problem as a blend between regularized multivariate regression and graphical models that combine the L_2,1_-norm with classical regularization techniques. We used data and networks from the DREAM5 challenge to show that the proposed models provide consistently good performance in comparison to contenders whose performance varies with data sets from simulation and experiments from model unicellular organisms *Escherichia coli* and *Saccharomyces cerevisiae*. Since the models’ formulation facilitates the prediction of master regulators, we also used the resulting findings to identify master regulators over all data sets as well as their plasticity across different environments. Our results demonstrate that the identified master regulators are in line with experimental evidence from the model bacterium *E. coli*. Together, our study demonstrates that simultaneous modelling of several target genes results in improved inference of GRNs and can be used as an alternative in different applications.

**Author summary:** Reconstruction of cellular networks based on snapshots of molecular profiles of the network components has been one of the key challenges in systems biology. In the context of reconstruction of gene regulatory networks (GRNs), this problem translates into inferring regulatory relationships between transcription factor coding genes and their targets based on, often small, number of expression profiles. While unsupervised nonlinear machine learning approaches have shown better performance than regularized linear regression approaches, the existing modeling strategies usually do predictions of regulators for one target gene at a time. Here, we ask if and to what extent the joint modeling of regulation for multiple targets leads to improvement of the accuracy of the inferred GRNs. To address this question, we proposed, implemented, and compared the performance of models cast as a blend between regularized multivariate regression and graphical models that combine the L_2,1_-norm with classical regularization techniques. Our results demonstrate that the proposed models, despite relying on linearity assumptions, show consistently good performance in comparison to existing, widely used alternatives.

## Introduction

Elucidation of gene-regulatory networks (GRNs), comprising the entirety of transcription factor (TF)-target gene interactions, remains one of the key challenges in systems biology studies of single cells and entire organisms [1]. Advances in technologies for probing gene-regulatory interactions, including: Chromatin immunoprecipitation combined with sequencing (ChIP-Seq) [2], Yeast one hybrid (Y1H) [3], and DNA-affinity purification sequencing (DAP-Seq) [4], have facilitated understandings in the *in vivo* and *in vitro* binding of TFs to the promoter region of target gene and have provided valuable resources for obtaining insights in the characteristics of GRNs across organisms [5, 6]. However, these technologies are still resource-intensive even when applied with model organisms. As a result, addressing this key challenge of systems biology necessitates the development of computational approaches for reconstruction of GRNs that rely on other data sources, such as gene expression, that capture in part the effect of TF binding and subsequent activation or repression of transcription of the target gene.

The computational approaches for GRN reconstruction use data from steady-state and/or time-resolved experiments; they rely on unsupervised, semi-supervised, and supervised machine learning methods [7] to identify TFs that explain the expression (patterns) of target genes (TGs). Recent advances in supervised learning of GRNs have benefited from the compendia of TF-target gene interactions obtained by the aforementioned technologies [8]. Irrespective of the data used and the machine learning approach applied, reconstruction of GRNs is often performed with considerably fewer observations (*n*) than number of predictors (*p*) that has resulted in the development and application of diverse regularization techniques in Gaussian graphical models (GGMs) [9, 10] and the regression setting [11, 12]. Further, due to the often non-linear dependence between the expression of TGs and their regulating TFs, machine learning techniques based on random forests [13–15] and kernels in combination with regressions [16] have resulted in improved accuracy of GRN reconstruction with data from *Escherichia Coli* and *Saccharomyces cerevisiae* [17].

Computational approaches for GRN reconstruction from gene expression data in the regression setting model the expression of each TG based on the expression of the TFs as predictors. In doing so, the relation between TGs is neglected in the process of model building [18]. Therefore, it remains unexplored if the simultaneous consideration of multiple TGs in the linear setting may perform as well as the models for individual targets in a non-linear setting.

Evidence from analysis of existing GRNs have demonstrated the presence of master regulators [19], *i*.*e*. TFs that regulate a sizeable proportion of target genes. The existing approaches either reconstruct GRNs assuming a prior that given TFs act as master regulators [20] or infer master regulators from the models built for the individual target genes. Furthermore, ChIPseq data have demonstrated the dependence of gene regulatory interactions on the biological context, determined by the interaction among the environment, developmental stage, and cell type/tissue [21]. Therefore, gene regulatory interactions are plastic and this characteristic is often neglected in the reconstruction of GRNs, particularly with data from multiple environmental perturbations and/or organisms, resulting in the reconstruction of consensus interactions [12].

To tackle these shortcomings, we propose two novel GRN reconstruction approaches as a blend between regularized multivariate regression and graphical models in the large-*p*-small-*n* setting. Specifically, by assuming that the observed gene expression data matrix is drawn from a multivariate normal distribution, we impose the L_2,1_-norm penalty on the regression coefficients along with the L_1_ (or L_2_) on the precision matrix to jointly model the gene expression of all TGs in the penalize likelihood framework. We then show that model formulation allows us to use an iterative scheme in which the estimate of the precision matrix is used to refine the regression coefficient estimates at the next iteration until convergence. Using gene expression data sets from *E. coli* and *S. cerevisiae* as well as *in silico* data from the Dialogue on Reverse Engineering Assessment and Methods (DREAM5) network inference challenges [17], we evaluate the performance of the proposed models via extensive comparative analyses with respect to the state-of-the-art methods and show the advantages of the proposed approaches in addressing the two mentioned shortcomings–the identification of master regulators and the detection of plastic interactions.

## Results and discussion

### Preliminaries and notation

Before presenting the models, which represents one of our results, we introduce the notation used in the rest of the manuscript. Let ***m***^***i***^ and ***m***_***j***_ be respectively the *i*^th^ row and *j*^th^ column of a matrix **M** = (*m*_*ij*_). **M**^−1^ and **M**^**T**^ represent respectively, the inverse and the transpose of **M. I**_*n*_ stands for the *n*-dimensional identity matrix, and if *m*_*i*_ is the *i*^th^ component of the vector ***m*** ∈ ℝ^*n*^, then its L_*p*_-norm is defined as

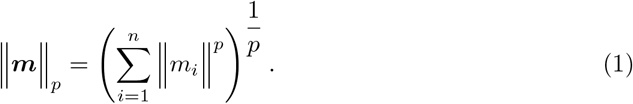

The L_2,1_-norm [36] of a matrix **M** ∈ ℝ^*k* × *l*^ and its partial derivative with respect to **M** are respectively

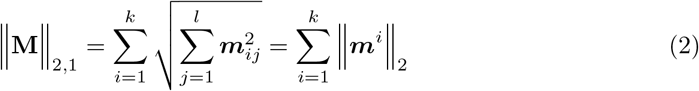

and 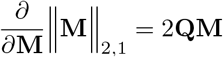, where **Q** ∈ ℝ^*k* × *k*^ is the diagonal matrix with entries 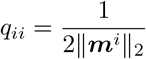.

In the regression setting for GRN inference, we aim to quantify the regulatory relationship between *s* TGs (i.e. response variables) ***y***_**1**_, *· · ·*, ***y***_***s***_ and a single set of *p* TFs (i.e. predictor variables) ***x***_**1**_, *· · ·*, ***x***_***p***_, such that ***y***_***k***_ = *b*_1*k*_***x***_**1**_ + *· · ·* + *b*_*pk*_***x***_***p***_ + ***ε***_***k***_, 1 ≤ *k* ≤ *s*. The model can then be cast in the matrix notation as

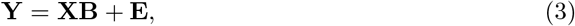

where **Y**_*n×s*_ = (***y***_**1**_, *· · ·*, ***y***_***n***_)^*T*^, **X**_*n×p*_ = (***x***_**1**_, · · ·, ***x***_***n***_)^*T*^, **B**_*p* × *s*_ = (***b***_**1**_, · · ·, ***b***_***p***_)^*T*^ and **E**_*n*×*s*_ = (***ε***_**1**_, · · ·, ***ε***_***n***_)^*T*^ are respectively the TGs (i.e. response), TFs (i.e. predictors), regulatory links (i.e. regression coefficients) and error matrices.

Assuming that the errors ***ε***_***i***_ are independent and normally distributed with covariance matrix 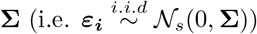, then the negative log-likelihood function [37] of the parameters (**B, Ω**) can be written up to a constant as

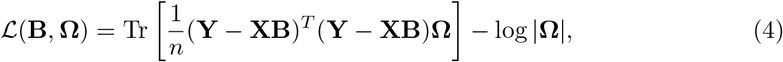

where **Ω** = **Σ**^−1^ is the precision matrix, Tr denotes the trace linear operator and |**Ω**| is the determinant of the matrix **Ω**. Estimators of the parameters **B** and **Ω** derived from standard procedures such as maximum likelihood and weighted least-squares are equivalent to those obtained when regressing each of the *s* responses on the *p* predictors separately. However, these estimators have poor performances, are computationally unstable and less efficient for prediction when the number of predictor and response variables are larger than the sample size.

As noted above, existing regression-based approaches for GRN reconstruction neglect the correlation among the response variables (*i*.*e*. TGs). To address this issue, we construct new sparse estimators for the regression coefficient and precision matrix via penalized likelihood optimization. Specifically, for tuning parameters *λ*_1_ ≥ 0, *λ*_2_ ≥ 0 and by penalizing the negative log-likelihood in Eq (4), the *s*(*s* + 1)*/*2 parameters of the precision matrix **Ω** are used to update the estimate of the regression coefficient **B** at the next iteration until convergence. In the following, we provide estimates 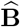 and 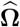 as solution to the mixed L_1_L_2,1_-norms and L_2_L_2,1_-norms regularized multivariate regression and covariance selection problems. For clarity, the terms experiment, condition and time point are used interchangeably; and mixed-norms terminology in this context simply refers to the fact that, the L_1_ (or L_2_) and L_2,1_ penalties are simultaneously imposed on **Ω** and **B** in the proposed optimization problems.

### Mixed L_1_L_2,1_-norms regularized multivariate regression and covariance selection

When the constant term with no effect on the optimization over **B** and **Ω** is ignored, the objective function to be minimized for the mixed L_1_L_2,1_-norms is proportional to

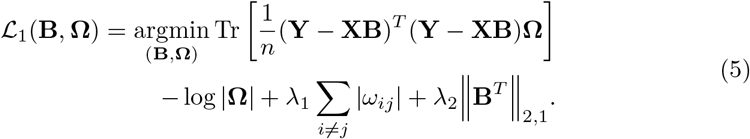

Notice how the L_2,1_ penalty is imposed on **B**^*T*^ instead of **B** ∈ ℝ^*p×s*^, since: (i) the number of TF genes (*p*) is generally considerably smaller than the number of TGs (*s*), and (ii) the L_2,1_ penalty may push some entries in **B** (i.e. TF-TG interaction) toward zero. As a result, this formulation facilitates model interpretation and the identification of candidate for interactions and master TFs.

The optimization problem in Eq (5) is biconvex. Therefore, convexity is ensured when solving for either parameter **B** or **Ω**, while keeping the other fixed. Solving for **B** with **Ω** fixed to **Ω**_0_, Eq (5) reduces to the convex:

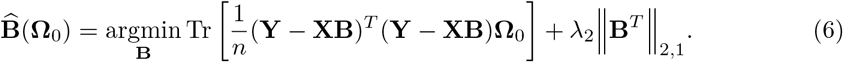

Taking the partial derivative with respect to **B** yields

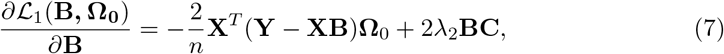

where **C** is the diagonal matrix with the *i*^th^ diagonal entry 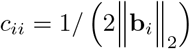. For computational stability, one can also use 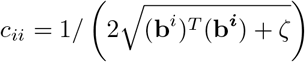 as an approximation [38], with *ζ* −→ 0.

#### Solving the mixed L_1_L_2,1_-norms model for B

The first-order condition defined by Eq (7) gives the following homogeneous Sylvester equation [39] in term of **B**:

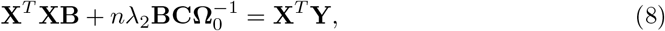

with solution

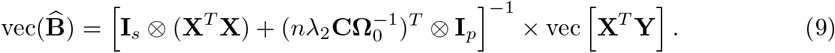

However, for gene expression data, *s* is often too large such that solution to Eq (8) as defined by Eq (9) becomes computationally prohibitive due to high memory requirements. We address this limitation (see Method 1 in S1 Appendix for details), by using the singular value decomposition (SVD) of 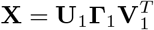, the matrix inversion lemma [40] and change of variables in Eq (10)

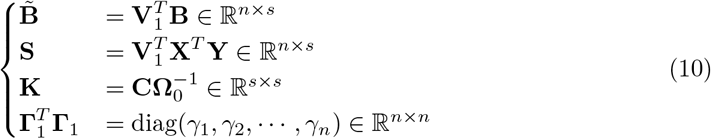

to obtain 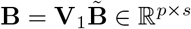. We refer to the latter as the L_1_L_2,1_-solution. Notice that, the proposed estimate can be viewed as a generalization of several existing approaches including: the multi-output regression [41] with identity task covariance (herein L_1_L_2,1_G-solution), the L_2,1_ feature selection [38], the ridge and ordinary least square. For detailed explanations regarding the derivation of these special estimates, we refer the reader to Method 2 in S1 Appendix.

#### Solving the mixed L_1_L_2,1_-norms model for Ω

For fixed **B** at a chosen point **B**_0_ and when solving for **Ω**, the optimization problem in Eq (5) yields

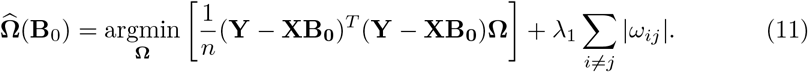

This corresponds to the L_1_-penalized covariance estimation problem and the graphical LASSO [42] (GLASSO) can be used to derive **Ω** for the model in Eq (11).

### Mixed L_2_L_2,1_-norms regularized multivariate regression and covariance selection

Analogously to the optimization problem in Eq (5), we formulate the following mixed L_2_L_2,1_-norms objective function:

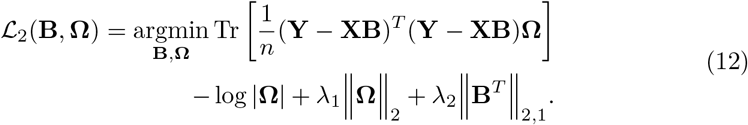

#### Solving the mixed L_2_L_2,1_-norms model for B

When solving for **B** with fixed **Ω**, the proposed mixed L_2_L_2,1_-norms model in Eq (12) which imposes the L_2_ penalty on **Ω** yields similar solutions as the optimization problem in Eq (6). Using similar methodology as S1 Method in S1 Appendix, we obtain the L_2_L_2,1_ and L_2_L_2,1_G-solutions, for respectively the main problem and the special case.

#### Solving the mixed L_2_L_2,1_-norms model for Ω

For fixed **B** at a chosen point **B**_0_ the optimization problem in Eq (12) when solving for **Ω** becomes

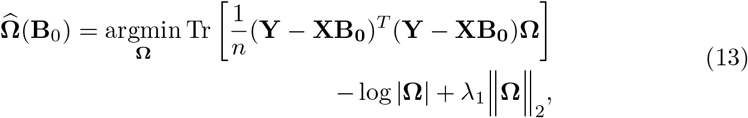

where the partial derivative with respect to **Ω** is given by

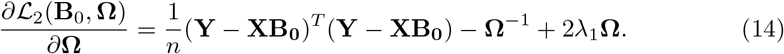

Defining 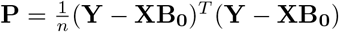 and setting Eq (14) to zero, we obtain the following quadratic matrix equation:

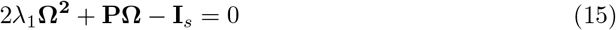

which is a special form of the well known algebraic Riccati equation encountered in multiple fields such as control theory and optimization [43, 44]. However, because the fundamental theorem of algebra is not valid for matrix polynomials, problems in the form of Eq (15) are often difficult to solve even in the matrix square root case **X**^2^ = **A** [45]. Therefore we ask if our problem then has a solution, which we answer by the affirmative (cf. Method 3 in S1 Appendix) and show that the solution to our problem exists and is uniquely given by

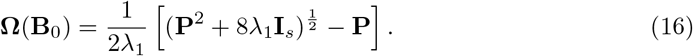

### Remark: Existence and uniqueness of a positive definite solution for quadratic matrix equations

It is known that equations of the form **AX**^**2**^ + **BX** + **C** = 0, **A, B, C** ∈ ∥^*s×s*^ can have no solution, a finite positive number or infinitely many solutions [22], but to the best of our knowledge, we found no particular evidence regarding the existence and uniqueness of solutions. However, while solving Eq (15) we noticed that, if **A** = **I**_*s*_, **B** and **C** commute and are respectively non-negative and positive definite, and if **B**^**2**^ − 4**C** is positive definite, then the existence and uniqueness of a positive definite solution **X** is guarantied and can be explicitly determined using the usual formula of the roots in the scalar case. With the positive definiteness requirement of the covariance and correlation matrices [9] being a major drawback in the situation where the sample size *n*, is smaller than the number of variables *s* (e.g. for microarray data sets), this existence and uniqueness of a positive definite solution can be of particular relevance when using GGM for GRN reverse engineering.

### Comparative analysis with DREAM5 data sets

The performance of the proposed inference approaches (i.e. L_1_L_2,1_, L_1_L_2,1_G, L_2_L_2,1_ and L_2_L_2,1_G) are compared with that of GENIE3, TIGRESS, ANOVerence, PLSNET, ENNET, PORTIA, etePORTIA and D3GRN when reconstructing the regulatory networks of *E. coli, S. cerevisiae* and the simulated data (i.e. *in silico*) with similar regulatory dynamic as *E. coli*. The contending methods are chosen to include the winner of the challenge (i.e. GENIE3, TIGRESS and ANOVerence), as well as some of the most recent state-of-the-art approaches (i.e. PLSNET, ENNET, PORTIA, etePORTIA and D3GRN) applied on the same data sets. For network-specific assessment and in contrast to all evaluated methods which exhibit large variability in performance across networks, Table 1 and S1 Fig, show that the proposed models show consistently good performance across all data sets. Overall and as depicted in the last three columns of Table 1 and S1 Table, the proposed approaches have comparable performances to that exhibited by the best method. Specifically, in terms of AUROC and Overall scores, the proposed models slightly outperform the contenders while the best performing state-of-the-art method (i.e. etePORTIA) in the comparative analysis shows an improved AUPR score of 1.3% compared to the former. With consistent performances across all evaluated data sets, we conclude that the proposed models are competitive and reliable alternatives to state-of-the-art GRN inference methods.

**Table 1.**
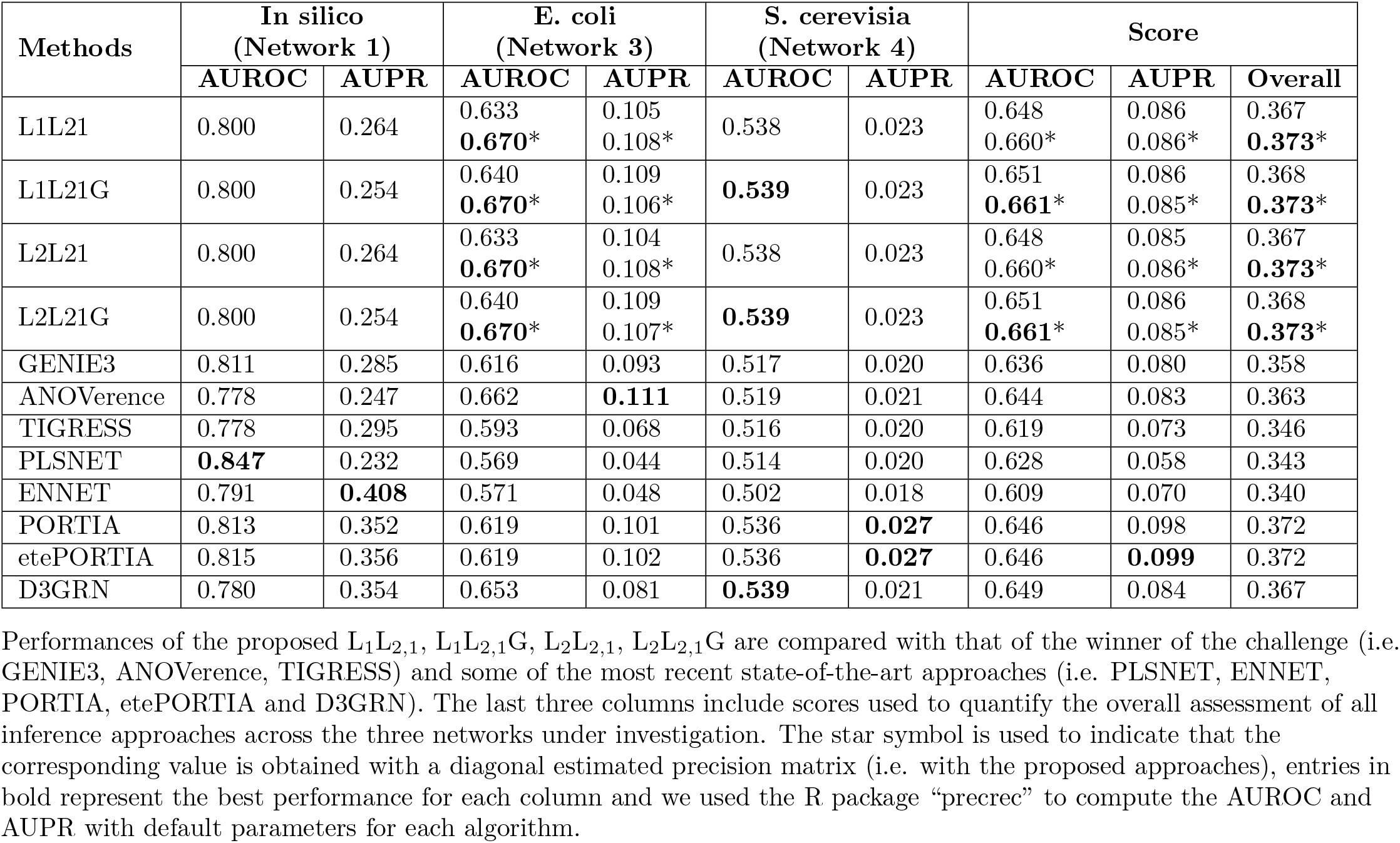
Comparison of model performance using area under the ROC curve (AUROC) and area under the precision-recall curve (AUPR) on DREAM5 data sets.

### Analysis with *E. coli* data across multiple conditions

Gene regulation depends on the cellular context including the cell type and the environmental conditions [23]. In this section, we focus on the latter and study master TFs involved in the regulatory dynamic of *E. coli* across multiple stress conditions. To this end, we applied our proposed models with data comprising few time-resolved samples gathered from *E. coli* strain MG1655 exposed to cold, heat, lactose-diauxic shift and oxidative stress conditions.

A recent comparative analysis with the same data contrasted the performance of the fused LASSO extension [12] with nine state-of-the-art inference methods. After re-evaluating all the models we reached a similar conclusion, whereby the fused LASSO achieves better performance and assigns higher scores to the true regulatory links. For this reason, we use the fused LASSO model as a benchmark when assessing the performance of our proposed approaches. Following the same methodology for performance assessment and for a fair comparison, a combination of TFs in RegulonDB [24] and DREAM5 challenge was considered, to finally obtain 173 TFs and 1561 TGs for GRN inference. Our findings summarized in Table 2 and (S2 Fig and S3 Fig) show that, except for the oxidative stress condition, the proposed inference methods outperform the fused LASSO (and contending approaches). As expected, and consistent with previous studies [25, 26], the results on combined data sets show an improved performance for all inference approaches with respect to AUROC and AUPR, as the sample size increased (i.e. from 5 for each condition to 20 for the combined data sets). Next, using the master regulator identification’s procedure (see Materials and methods) and considering all TFs interacting with more than 50% (i.e. *α*) TGs, we compiled in Table 3A, the list of MR^1^ conserved across all conditions. Although originally designed for gene tissue specificity, we adapted the *τ* -index [27] as shown in Eq (17) to compute condition specificity of MR^1^ and MR^2^ that we previously identified to be conserved across conditions

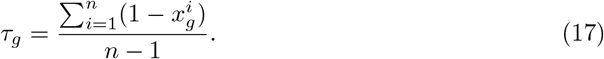

**Table 2.** Comparative analysis based on time-resolved transcriptomics data sets for model organism *E. coli*.

| Methods | Cold |  | Heat |  | Oxidative |  | Lactose |  | Combined data sets |  | Score |  |  |
| --- | --- | --- | --- | --- | --- | --- | --- | --- | --- | --- | --- | --- | --- |
|  | AUROC | AUPR | AUROC | AUPR | AUROC | AUPR | AUROC | AUPR | AUROC | AUPR | AUROC | AUPR | Overall |
| L1L21 | 0.526 | **0.015** | 0.515 | **0.014** | 0.521 | **0.015** | 0.501 | 0.013 | **0.564** | **0.017** | 0.524 | **0.014** | 0.269 |
| L1L21G | **0.535** | **0.015** | **0.516** | **0.014** | 0.502 | 0.013 | **0.504** | **0.014** | 0.561 | **0.017** | 0.523 | **0.014** | 0.268 |
| L2L21 | 0.529 | **0.015** | 0.515 | **0.014** | 0.521 | **0.015** | 0.501 | 0.013 | **0.564** | **0.017** | **0.525** | **0.014** | **0.270** |
| L2L21G | 0.530 | **0.015** | 0.509 | **0.014** | 0.502 | 0.013 | **0.504** | **0.014** | 0.561 | **0.017** | 0.520 | **0.014** | 0.267 |
| Fused LASSO | 0.494 | 0.013 | 0.480 | 0.012 | **0.532** | **0.015** | 0.502 | 0.013 | 0.539 | **0.017** | 0.508 | 0.013 | 0.261 |

The performances of the proposed methods are contrasted with that of Fused LASSO under four experimental conditions including heat, cold, lactose, oxidative as well as their combination. Scores in the last three columns are also shown to quantify the overall performance of the inference approaches across all data sets. Entries in bold represent the best performance and we used the R package “precrec” to compute the AUROC and AUPR.

**Table 3.**
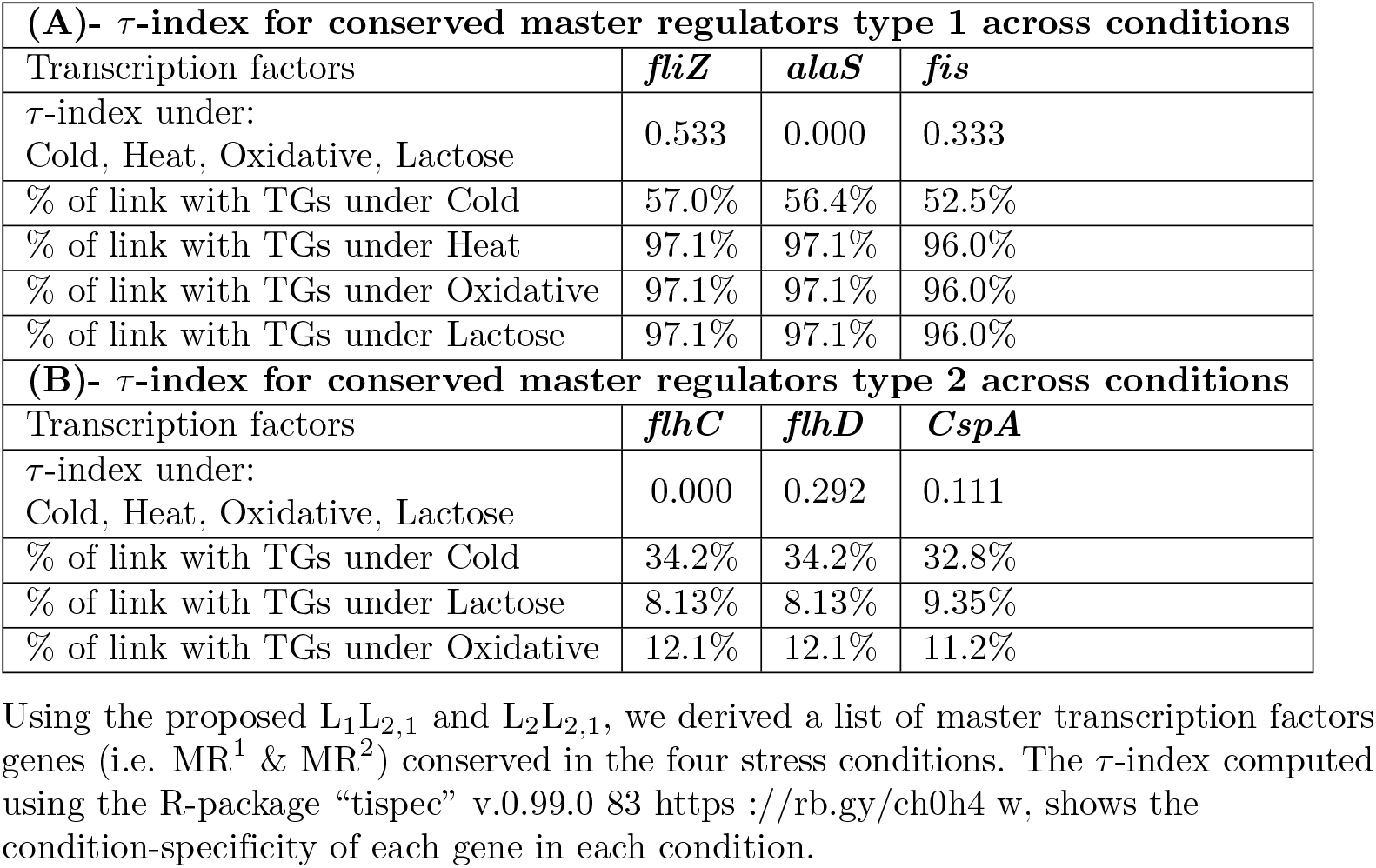
Specificity index for MR^1^ & MR^2^ across cold, heat, lactose and oxidative conditions.

Here, the number of conditions is denoted by *n* and 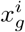 represents the mean expression profile of gene *g* in the *i*^th^ condition. It can be observed that *τ* ∈ [0, 1] and depending on the obtained value, we infer that the corresponding master TF is a housekeeping gene (i.e. *τ* → 0) or condition-specific (i.e. *τ* → 1). Following [27] and [28], that respectively considered *τ* ≥ .85 and .8 as a threshold for tissue specificity, we use as decision rule (*τ* > .8) to check if the given master regulator is ubiquitously expressed or not. Interestingly our finding is in agreement with the *τ* -index (cf. Table 3A), whereby all MR^1^ that the proposed L_1_L_2,1_ and L_2_L_2,1_ found conserved across all four conditions have a maximum condition specificity index of .533. Using the derived *τ* -index as a sanity check, we conclude that these master transcription factors are indeed conserved across all conditions. In contrast, MR^2^ are only found conserved across three of the four stress conditions (i.e. cold, lactose and oxidative). This is in line with the study by [29] in which it was suggested that *E. coli* perceives high temperatures as a sign of inflammation, and as a result downregulates flagella class II and III genes (to avoid detection by the host immune system). This process is caused by the lower level of upstream activator *flhD* that we found conserved under other three stress conditions. Additionally, the presence of *flhD* and *flhC* in our list of conserved master regulator is quite interesting as these have been previously identified as master regulator for the expression of flagellar genes in *E. coli* [30, 31]. Similarly, the absence of conservation of the transcription factor *CspA* under heat condition could be justified, since it is among the major cold shock proteins of *E. coli* [32] that are only induced upon temperature decrease. Specifically, it has been shown that the induction of *CspA* is mainly caused by dramatic stabilization of its mRNA at low temperature [33, 34].

The study of sparsity level in our estimated regression coefficients and precision matrix shows that the expression of 1,156 genes was under the regulation of all 173 TFs used for the analysis (i.e. none of the rows of regression coefficients or in the precision matrix was entirely zero). Cold was the stress condition for which the three MR^1^, *fliZ, alaS* and *fis*, regulated respectively about 57%, 56% and 53% of the 1,156 genes (cf. Table 3A). In contrast, lactose was the stress for which the MR^2^ regulated the smallest numner of TGs (cf. Table 3B). To further investigate if the conserved MR^1^ and MR^2^ share any biological attributes, we performed enrichment analysis using the web application “ShinyGO” [35] while correcting for multiple testing with false discovery rate (FDR) (p-value *<* 0.05). The enrichment analysis (GO biological process) reveals that the conserved MR^1^ (cf. S4 FigA) are mostly enriched for negative regulation of RNA biosynthesis process, nucleic acid-templated transcription and nucleobase-containing compound metabolic process. Moreover, MR^2^ (cf. S4 FigB) conserved under cold, lactose and oxidative stress conditions are mostly enriched in three biological processes including regulation of organelle, bacterial-type flagellum and cell projection assembly.

## Conclusion

We proposed two novel approaches that cast the GRN reconstruction problem as a blend between regularized multivariate regression and graphical models. Through extensive comparative analysis with simulated and real-world data, we demonstrated that the introduced models are consistent and exhibit excellent performance over the contenders. Considering the often encountered dilemma in GRN inference whereby a choice has to be made between linear and non-linear modeling assumptions, we further show that consideration of multiple responses even in a linear setting can show as good performance as non-linear approaches (e.g. random forests). In addition, without assuming any prior on TFs nor inferring them from the individual models built for the target genes, the L_1_L_2,1_ and L_2_L_2,1_ leverage sparsity in the regression coefficients and precision matrix to identify master regulators while offering the possibility to infer their plasticity and regulatory interactions.

## Materials and methods

### Data sets

#### DREAM5

To evaluate the performance of the proposed and contending approaches, we use three benchmark data sets from the DREAM5 challenge freely available from [17]. As summarized in Table 4, each data set contains a collection of gene expression profiles, a gold standard (*i*.*e*. a set of verified interactions) and a list of known TFs.

**Table 4.** Details of gene expression data sets for model organisms *E. coli, S. cerevisiae*, as well as *in silico* from DREAM5.

| Networks | #Samples | #TFs | #Genes | #Verified interactions |
| --- | --- | --- | --- | --- |
| *In silico* (Network 1) | 805 | 195 | 1643 | 4012 |
| *E. coli* (Network 3) | 805 | 334 | 4511 | 2066 |
| *S. cerevisiae* (Network 4) | 536 | 333 | 5950 | 3940 |

For each network, this includes the number of putative TFs, TGs, samples and verified interactions in the gold standard. The original labels of each network from the challenge are given in parentheses.

Briefly, network 1 is a simulated data set mimicking the transcriptional regulatory network of *E. coli* in which 10% of random edges were added and the expression profile generated with GeneNetWeaver [53]. For network 3 and network 4, the Gene Expression Omnibus (GEO) database [54] was used to produce affymetrix genuine gene expression data sets for *E. coli* and *S. cerevisiae* respectively. The resulting microarray data sets where then normalized using Robust Multichip Averaging (RMA) [55]. For a detailed description of the DREAM5 inference challenge, its design and the data generation process, interested readers are referred to [17] and the DREAM website.

#### *E. coli* time-resolved transcriptomics data

The ability of the proposed methods to reconstruct GRN with small sample data across multiple conditions or tissues is evaluated by further considering time-resolved transcriptomics data resulting from the experiment in [56], available from the GEO database under accession GSE20305. Here, we investigated the gene expression responses of *E. coli* strain MG1655 to four stress conditions (*i*.*e*. oxidative stress, glucose-lactose diauxic shift, heat, and cold). Except for the scenario where stress was induced by hydrogen peroxide (*i*.*e*. oxidative stress), sampling with 10 min steps for transcript profiling was performed from time points 10-50 min post-perturbation plus two control time points prior to each perturbation. Averaging over the three available biological replicates for each time point resulted to the expression profile data of five samples for individual stress condition and 4400 genes.

### Data pre-processing, hyperparameter tuning and evaluation metrics

As a pre-processing step, the expression levels of each gene are centered and scaled within each data set. To tune hyperparameters *λ*_1_ and *λ*_2_, we used 10-fold cross-validation (CV) and split each gene expression profile data set from DREAM5 into 10 non-overlapping subsets of almost identical size. With 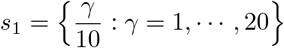 and *s*_2_ = {2^−*δ*^ : *δ* = 0, *· · ·*, 8} as the search spaces for *λ*_1_ and *λ*_2_ respectively, we finally select the optimal *λ*_1_ and *λ*_2_ as the maximizer of the log-likelihood on the validation data. Due to the very small sample size in the case of time-resolved data sets, leave-one-out CV was used instead with the same grids. Interestingly, we observed that model performance is more influenced by *λ*_2_, the penalty on the regression coefficient matrix. We further found that there is a limiting factor for which irrespective of the chosen *λ*_1_, **Ω** results in a diagonal matrix. This is very useful for the practical implementation as it can be used to efficiently reduce computation time while controlling the amount of sparsity in the precision matrix.

Regarding performance evaluation, we follow the DREAM5 strategy and only consider the top 100,000 edge predictions to evaluate TF-TG interactions as a binary classification problem for which, edges are predicted to be present or absent. With the selected interactions, we then make use of area under the receiver operating characteristic (AUROC) and area under the precision-recall (AUPR) curves, two widely used metrics for performance assessment in GRN inference. For an overview of the performances across all used data sets, we also computed the score for each metric and the overall score as shown in Eq (18).

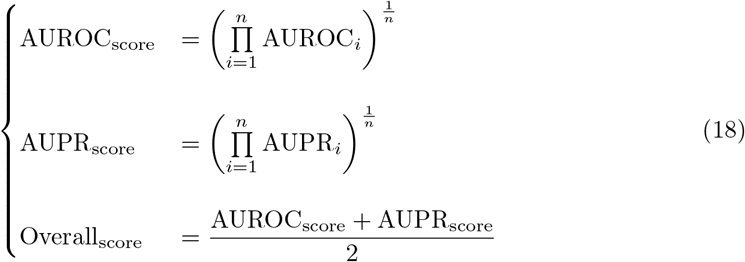

where *n* is the number of considered networks (*e*.*g*. in the current analysis, *n* = 3 and *n* = 5 for respectively DREAM5 and time-resolved transcritomics data sets). In addition, recalling that for the proposed approaches we would like to quantify the contribution of individual TF on the remaining genes (*i*.*e*. respectively rows and columns of our estimated regression coefficient matrices), we scale TF-wise, edge weights obtained from each inference method to range in the interval [0, 1]. That is, for the *i*^th^ row ***β***^*i*^ = [*β*_1_, *· · ·, β*_*s*_] of the estimated coefficient matrix, the maximum absolute scaling is used to compute each normalized entry as 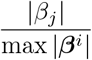.

### Contending approaches

To provide a comprehensive comparative analysis, we compared the solutions of the proposed models with nine state-of-the-art approaches. To account for updated developments in GRN inference and because our analysis relies on the data sets from DREAM5 challenge, we selected D3GRN [46], PLSNET [47], ENNET [48], PORTIA and its extension etePORTIA [49] as some of the most recent state-of-the-art approaches that used the same data sets. Further, we included those methods that were ranked among the top three GRN reconstruction approaches in the challenge based on the overall score. These approaches included: TIGRESS [14], that was deemed the best linear regression-based method in DREAM5, GENIE3 [50], that uses variable selection with ensembles of regression trees and ANOVerence [51] that relies on the non-linear Cohen’s correlation coefficient *η*^2^ computed from two-way analysis of variance (ANOVA). We also included the Fused LASSO [12] formulation that combines information from multiple data sets, shown to outperform contending approaches.

### Identification of master TFs

The term “master regulator” refers to a TF that is at the top of the transcriptome regulatory hierarchy, thus regulating the majority of other TFs and associated TGs [52]. We use the estimated sparse regression coefficient and precision matrix from the proposed models to identify the master regulator type 1 and type 2 (i.e.MR^1^ and MR^2^). Given the estimated sparse regression coefficient matrix 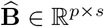 and precision matrix 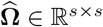, we say that a TF (i.e. column of the predictor matrix **X** ∈ ℝ^*n×p*^) is a type 1, *α* − master regulator 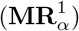 if for 0 *< α* ≤ 1, the corresponding row in 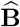 has an *α* − percentage of non-zero entries. Regarding type 2 master regulator 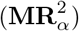, we use conditional dependence (i.e. non-zero TF-TG entries in the sparse precision matrix) to validate that the same TF-TG in 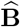 is non-zero. While enhancing the sparsity in the regression coefficient matrix, this procedure also serves to validate if the direct link identified by 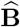 remains a link given the rest of genes in the network. Finally, similar to type 1, 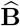 derived from this procedure is then used to detect what we call 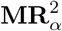.

## Availability and implementation

The approaches are implemented using the R programming language and the codes are freely available from https://github.com/alainmbebi/mixed-norms-GRN. All data underlying this publication are publicly available and their corresponding references provided in the article.

## Funding

AJM & ZN are supported by the European Union’s Horizon 2020 research and innovation programme in connection with the projects BREEDCAFS [GA No. 727934] https://www.breedcafs.eu/ and PlantaSYST [FPA No. 664620] https://plantasyst.eu/. The funders had no role in study design, data collection and analysis, decision to publish, or preparation of the manuscript.

## Competing interests

The authors have declared that no competing interests exist.

## Supporting information

**S1 Fig. PR and ROC curves for individual methods in the comparative analysis with DREAM5 data sets**.

We used the following models: the L_1_L_2,1_, L_1_L_2,1_G, L_2_L_2,1_, L_2_L_2,1_G, the winner of the challenge (i.e. GENIE3, ANOVerence, TIGRESS) and some of the most recent state-of-the-art approaches (i.e. PLSNET, ENNET, PORTIA, etePORTIA and D3GRN) to infer the regulatory networks of *E. coli* (left), *S. cerevisiae* (middle) and *in silico* (right). Shown in the upper and lower panels are respectively the precision-recall (PR) and receiver operating characteristic (ROC) curves.

**S2 Fig. PR curves for individual methods in the comparative analysis with *E. coli* time-resolved transcriptomics data sets**.

We used the following models: the L_1_L_2,1_, L_1_L_2,1_G, L_2_L_2,1_, L_2_L_2,1_G and Fused LASSO to infer the regulatory networks of *E. coli* from time-resolved transcriptomics experiment. Shown are the precision-recall (PR) curves for each inference methods under heat, cold, lactose, oxidative conditions and when all data sets are combined.

Recalling that with the same data sets, the fused LASSO was already assessed and outperformed the contending approaches, the current comparative analysis implicitly extends to Gaussian graphical models (GGM), the algorithm for the reconstruction of accurate cellular networks(ARACNE), GENIE3, global silencing, CLR and LASSO-type (i.e. L_1_, L_0_ and L_1*/*2_) regularization.

**S3 Fig. ROC curves for individual methods in the comparative analysis with *E. coli* time-resolved transcriptomics data sets**.

We used the following models: the L_1_L_2,1_, L_1_L_2,1_G, L_2_L_2,1_, L_2_L_2,1_G and Fused LASSO to infer the regulatory networks of *E. coli* from time-resolved transcriptomics experiment. Shown are the receiver operating characteristic (ROC) curves for each inference methods under heat, cold, lactose, oxidative conditions and when all data sets are combined. Recalling that with the same data sets, the fused LASSO was already assessed and outperformed the contending approaches, the current comparative analysis implicitly extends to Gaussian graphical models (GGM), the algorithm for the reconstruction of accurate cellular networks (ARACNE), GENIE3, global silencing, CLR and LASSO-type (i.e. L_1_, L_0_ and L_1*/*2_) regularization.

**S4 Fig. Enrichment analysis of conserved MR**^1^ **and MR**^2^ **with time-resolved transcriptomic data sets from *E. coli***.

Shown are the fold enrichment sorted by GO biological process. (A) MR^1^ found conserved across the four stress conditions. (B) MR^2^ conserved under cold, lactose and oxidative stress. We used the graphical gene-set enrichment tool “ShinyGO” v.0.76.1 http://bioinformatics.sdstate.edu/go/ for the analysis.

**S1 Table. Comparison of model performance using area under the ROC curve (AUROC) and area under the precision-recall curve (AUPR) on DREAM5 data sets**.

The reported results are from the DREAM5 challenge and correspond to the best (i.e. overall score) inference methods that participated in the challenge. Since results obtained using the R package “precrec” were slightly different from those of the challenge (cf. Table 1), we sought to include the latter here to have a comprehensive assessment and to avoid misinterpretation of the current results.

## S1 Appendix

### Supplementary methods

The appendix includes detailed explanations on how to derive: (1) The matrix of regression coefficients B, as the solution to a special case of Sylvester equation, (2) The special cases of the L_1_L_2,1_ and L_2_L_2,1_ solutions as well as the precision matrix Ω as the solution to a special form of algebraic Riccati equation.

## Notes

### Competing Interest Statement

The authors have declared no competing interest.

